# Chemogenetic attenuation of PFC pyramidal neurons restores deficits in recognition memory following adolescent NMDA receptor blockade

**DOI:** 10.1101/2024.01.10.574791

**Authors:** Hagar Bauminger, Sailendrakumar Kolatt Chandran, Irit Akirav, Inna Gaisler-Salomon

## Abstract

During adolescence, the prefrontal cortex (PFC) undergoes dramatic developmental changes, including fine-tuning the balance between excitatory glutamate and inhibitory GABA transmission (i.e., the E/I balance). This process is critical for intact cognitive function and social behavior in adulthood, and its disruption is associated with several psychiatric disorders including schizophrenia (SZ). While acute NMDA receptor (NMDAr) blockade leads to excess glutamate transmission in the PFC, the long-term consequences of MK-801 administration during early adolescence on the E/I balance in adulthood have not been extensively studied. In the current study, we show that chronic MK-801 administration during early adolescence leads to abnormalities in recognition memory and social behavior as well as reduced frequency of miniature inhibitory post-synaptic currents (mIPSCs) in mPFC of adult male rats, with no change in excitatory currents or basal activity. We further show that chemogenetic attenuation of prelimbic mPFC pyramidal neurons reversed deficits in recognition memory, but not social behavior. These findings emphasize the critical role played by NMDAr during adolescence on the E/I balance as well as cognition and social function in adulthood. Moreover, these findings implicate the therapeutic outcomes of reduced mPFC pyramidal neuron activity in recognition memory deficits in early-adolescence MK-801-treated rats. Since recognition memory deficits are key components of the cognitive deficits in SZ, these findings suggest that the future development of treatments aimed at alleviating the cognitive deficits in SZ should focus on regulating the prefrontal E/I balance.

## Introduction

Adolescence is a vulnerable developmental time period during which several psychiatric disorders emerge. The prefrontal cortex (PFC) undergoes dramatic developmental changes during adolescence (Blakemore, 2012b). In particular, intact PFC development during the peri-pubertal years is critical for cognitive function, e.g., memory and problem-solving abilities (Bauminger & Gaisler-Salomon, 2022), and for social capacities (Blakemore, 2012a). Both of which are disrupted in schizophrenia (SZ; Lewis and González-Burgos, 2008). Preclinical studies point to the medial PFC (mPFC), equivalent to the human dorsolateral PFC as a key mediator of social and cognitive function in rodents (Uylings, Groenewegen, & Kolb, 2003).

A closer examination of the synaptic processes in the developing PFC indicates that adolescence is characterized by fine-tuning of the balance between excitatory (glutamatergic) and inhibitory (GABAergic) activity (i.e., the E/I balance; Insel, 2010). This balance is achieved by increasing inhibitory and decreasing excitatory synaptic strength in the cortex during adolescence (Caballero, Granberg, & Tseng, 2016; Insel, 2010). Aberrant E/I balance was hypothesized to be a core feature of several neurodevelopmental disorders characterized by social and cognitive deficits, including SZ (Lopatina et al., 2019).

Non-competitive NMDA receptor (NMDAr) antagonists (e.g., MK-801, ketamine, PCP) are commonly used in pre-clinical studies to examine the neural basis of SZ-like cognitive deficits and negative symptoms (Neill et al., 2010). Previous studies on acute NMDAr blockade have pointed to enhanced cortical glutamate release (Moghaddam, Adams, Verma, & Daly, 1997) and changes in pyramidal neuron firing patterns (Homayoun & Moghaddam, 2007; Wang et al., 2013). Few studies have investigated the long-term effects of chronic NMDAr blockade, particularly during adolescence. In a previous study, we showed that when administered chronically in adolescence, NMDAr blockade in rats leads to recognition memory impairments and social interaction deficits in the adult animal. In parallel, we observed changes in the expression of glutamate and GABA markers, suggesting a dysregulated E/I balance (Bauminger, Zaidan, Akirav, & Gaisler-Salomon, 2022). Despite the centrality of adolescence for maturation of the E/I balance and relevance for cognitive and social functions, less attention has focused on examining the long-term impact of chronic NMDAr blockade on excitatory and inhibitory transmission in mPFC. Recently, MK-801 administration in adolescent male rats was found to reduce miniature inhibitory postsynaptic currents (mIPSCs) in pre-limbic (PrL) layer V (LV) pyramidal cells (Flores-Barrera, Thomases, & Tseng, 2020). However, the relevance of pertubed E/I balance following adolescent MK-801 exposure to cognitive and social behavior abnormalities has not been tested. Moreover, the restorative ability of PrL pyramidal neuron attenuation on adolescence MK-801-induced cognitive and social behavioral deficits has not been previously assesed.

Here, we first conducted a slice physiology study to assess the long-term impact of chronic MK-801 administration in early adolescence on the basal and synaptic properties of pyramidal neurons in the adult mPFC. Next, we tested whether chemogenetic attenuation of mPFC pyramidal neurons can reverse the early-adolescence MK-801-induced deficits in recognition memory and social behavior.

## Materials and Methods

### Subjects

Male Sprague Dawley rats (25 days old, weighing 70-100g; Envigo Laboratories), were group-housed (4 animals per cage) at 22±2°C under 12hr light/dark cycles. Animals had ad libitum access to food and water. Experimental procedures were approved by the University of Haifa Ethics and Animal Care Committee (approval number UoH-IL-2112-115).

### Pharmacological agents

The noncompetitive NMDAr antagonist MK-801 (dizocilpine; 0.2 mg/kg; Sigma Aldrich, Israel) was dissolved in 0.9% saline. Controls received saline. Clozapine n-oxide (CNO; 1mg/kg, Enzo) was dissolved in 0.9% saline. MK-801 and CNO were injected intraperitoneally (i.p.) at 1 ml/kg. Drug doses were based on previous publications (Bauminger et al., 2022; Maestas-Olguin, Fennelly, & Pentkowski, 2021).

### Experimental design

Figure 1 summarizes the experimental design. In Experiment 1, the long-term electrophysiological consequences of early-adolescence MK-801 or saline administration on LV PrL pyramidal cells were tested. Rats received chronic i.p. injections of saline or MK-801 for 15 days starting in early adolescence (Postnatal days (PND) 30-44), and electrophysiological recordings were performed in adulthood (between PND 60-85). In Experiment 2, chemogenetic attenuation of PrL pyramidal cells activity on MK-801-induced SZ-like behavioral deficits was examined. MK-801 injections were performed as in Experiment 1, and on PND60-68 rats underwent stereotactic surgeries to bilaterally inject viral vectors expressing pAAV-CaMKІІa-hM4D(Gi)-mCherry (i.e., hM4Di) or the control vector pAAV-CaMKІІa-mCherry (i.e., control) into the PrL. Behavioral tests were performed between PND90-98. The behavioral battery consisted of an open field (OF) test of locomotor activity and anxiogenic behavior, a novel object recognition (NOR) task, and a social interaction test (SIT), conducted 5min after the NOR test phase (Bauminger et al., 2022). CNO was administered 30min before the NOR task. A week later, rats were anesthetized, and brains were perfused; post-fixation brains were kept at −80°C.

**Figure 1.**
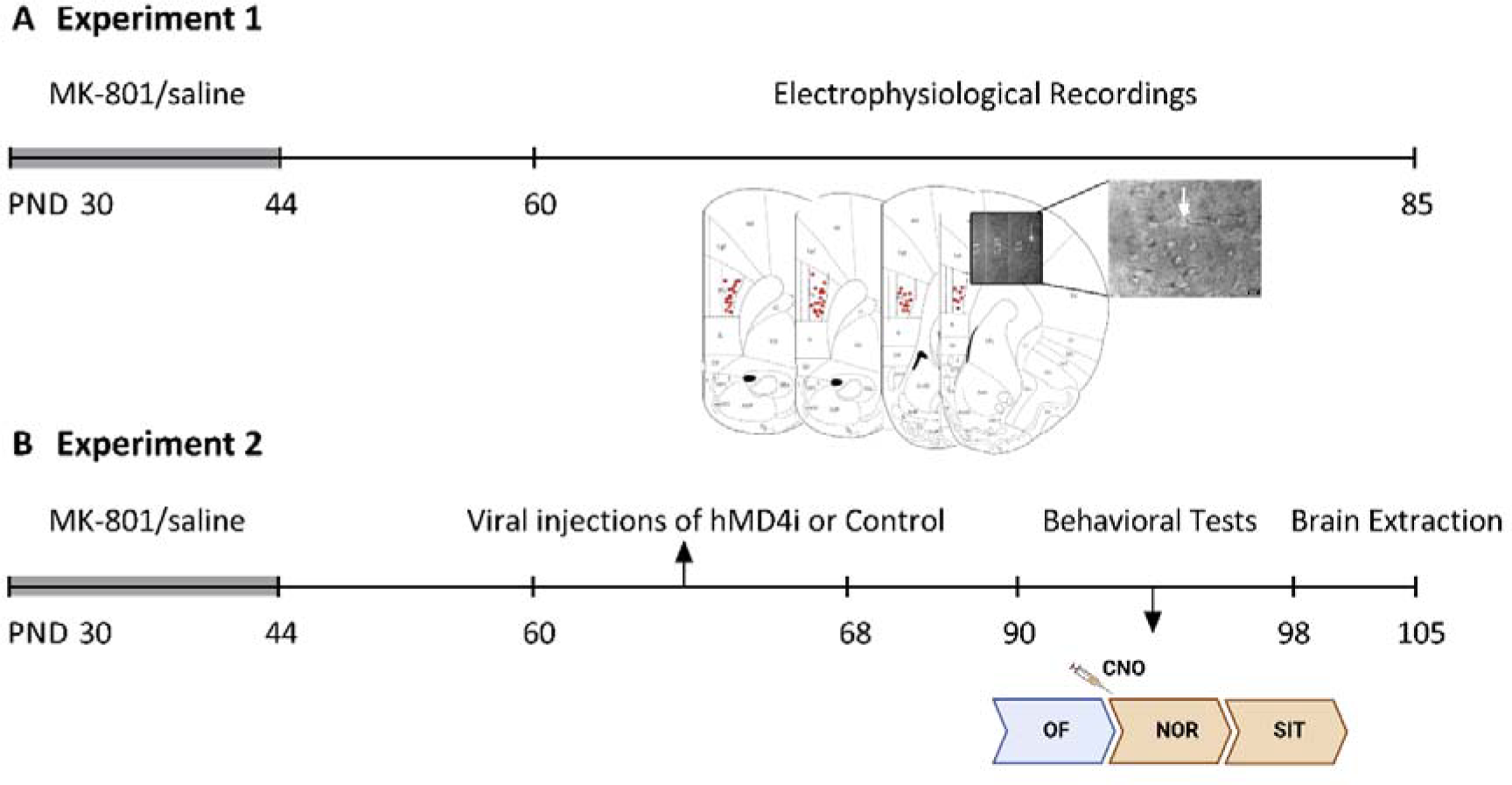
Experimental procedure timeline. (A) In Experiment 1, rats were administered MK-801 or saline in early adolescence, and electrophysiological assessment of PrL pyramidal cells was performed in adulthood as can be seen in the representative image showing the patch pipette positions (insets) and recorded areas (represented in red dots). (B) In Experiment 2, rats received systemic MK-801 or saline in early adolescence, and in adulthood intra-PrL injections of hM4Di or control virus under the controlled expression of CaMKІІa promoter, were performed. Behavior in the OF and CNO-induced behavioral changes in the NOR task and SIT were assessed 4 weeks after viral injections. CNO= Clozapine N-oxide, NOR=Novel Object Recognition, OF=Open Field, PND=Postnatal day, SIT=Social Interaction Test.

### Slice electrophysiology and whole-cell patch clamp

The slice electrophysiology and recording parameters were used as described previously (Chandran et al., 2022; Yiannakas et al., 2021). Briefly, rats were deeply anesthetized using isoflurane, and brains were extracted following decapitation. Three hundred µm thick coronal brain slices were obtained using a Campden-1000® Vibratome. Slices were cut in an ice-cold sucrose-based cutting solution containing the following (in mM): 110 sucrose, 60 NaCl, 3 KCl, 1.25 NaH2PO4, 28 NaHCO3, 0.5 CaCl2, 7 MgCl2, 5 D-glucose, and 0.6 ascorbate. Slices were allowed to recover for 30 min at 37°C in artificial CSF (ACSF) containing the following (in mM): 125 NaCl, 2.5 KCl, 1.25 NaH2PO4, 25 NaHCO3, 25 D-glucose, 2 CaCl2, and 1 MgCl2. Slices were kept for an additional 30 min in ACSF at room temperature until electrophysiological recording began. The solutions were constantly infused with carbogen (95% O2, 5% CO2).

### Intracellular whole-cell recording

After a recovery period, slices were placed in the recording chamber and maintained at 32-34°C with continuous perfusion of carbonated ACSF (2 ml/min). Brain slices containing the PrL cortex were illuminated with infrared light and pyramidal cells were visualized under a differential interference contrast microscope with 10X or 40X water-immersion objectives mounted on a fixed-stage microscope (BX51-WI; Olympus®). The image was displayed on a video monitor using a charge-coupled device (CCD) camera (Orca Retiga 2®, Hamamatsu Japan). Recordings were amplified by Double IPA® Integrated Patch Clamp Amplifiers with Data Acquisition System (Sutter Instruments®). The recording electrode was pulled from a borosilicate glass pipette (3–5 M) using an electrode puller (P-1000; Sutter Instruments®) and filled with a K-gluconate-based internal solution containing the following (in mM): 130 K-gluconate, 5 KCl, 10 HEPES, 2.5 MgCl2, 0.6 EGTA, 4 Mg-ATP, 0.4 Na3GTP and 10 phosphocreatine (Na salt). The osmolarity was 290 mOsm, and the pH was 7.3.

The recordings were made from the soma of LV pyramidal cells in the PrL cortex. Liquid junction potential (10 mV) was not corrected online. All current clamp recordings were low pass filtered at 10 kHz and sampled at 50 kHz. Pipette capacitance and series resistance were compensated and only cells with series resistance smaller than 20 MΩ were included in the dataset. Data quantification was done with Sutter Patch (Version 2.2, Sutter Instruments, Novato, CA) and Clampfit (Molecular Devices, Sunnyvale, CA) and subsequently analyzed using GraphPad Prism®. The method for measuring active intrinsic properties was based on a modified version of previous protocols (Chakraborty, Fedorova, Bagrov, & Kaphzan, 2017; Kaphzan et al., 2013; Sharma et al., 2017).

### Recording parameters

Intrinsic properties: Resting membrane potential (RMP) was measured 10 seconds after the beginning of the whole cell recording (rupture of the membrane under the recording pipette). The dependence of the firing rate on the injected current was obtained by injection of current steps (of 500ms duration from 0 to 400 pA in 50 pA increments). Input resistance was calculated from the voltage response to a hyperpolarizing current pulse (−150 pA). SAG ratio was calculated from voltage response −150 pA. The SAG ratio during the hyperpolarizing steps was calculated as [(1-ΔVSS/ ΔV max) x 100%] as previously reported by (Song, Ehlers, & Moyer, 2015). The membrane time constant was determined using a single exponential fit in the first 100ms of the raising phase of cell response to a 1 second, −150 pA hyperpolarization step.

For measurements of a single action potential (AP), after an initial assessment of the current required to induce an AP at 15ms from the start of the current injection with large steps (50 pA), a series of brief depolarizing currents were injected for 10ms in steps of 10 pA increments. The first AP that appeared at the 5ms time point was analyzed. A curve of dV/dt was created for that trace and the 30 V/s point in the rising slope of the AP was considered as threshold (Chakraborty et al., 2017). AP amplitude was measured from the equipotential point of the threshold to the spike peak, whereas AP duration was measured at the point of half amplitude of the spike. The medium after-hyperpolarization (mAHP) was measured using prolonged (3 seconds), high-amplitude (3 nA) somatic current injections to initiate time-locked AP trains of 50 Hz frequency and duration (10 –50 Hz, 1 or 3 s) in pyramidal cells. These AP trains generated prolonged (20 s) AHPs, the amplitudes, and integrals of which increased with the number of APs in the spike train. AHP was measured from the equipotential point of the threshold to the anti-peak of the same spike (Gulledge et al., 2013). Fast (fAHP), and slow AHP (sAHP) measurements were identified as previously described (Andrade, Foehring, & Tzingounis, 2012; Song & Moyer, 2018). Series resistance, Rin, and membrane capacitance were monitored during the entire experiment. Changes of at least 30% in these parameters were criteria for the exclusion of data.

Synaptic properties: Both miniature inhibitory and miniature excitatory postsynaptic currents (mIPSCs and mEPSCs, respectively) were measured following the application of antagonists in a perfusing ASCF containing cocktails of AMPA, NMDA, GABA, and TTX. To record mEPSCs, the recording electrode was pulled from a borosilicate glass pipette (3–5 M) using an electrode puller (P-1000; Sutter Instruments®) and filled with a K-gluconate-based internal solution containing the following (in mM): 130 K-gluconate, 5 KCl, 10 HEPES, 2.5 MgCl2, 0.6 EGTA, 4 Mg-ATP, 0.4 Na3GTP and 10 phosphocreatine (Na salt). The osmolarity was 290 mOsm, and pH of 7.3. mEPSCs were recorded in voltage clamp mode at a holding potential of −70 mV. 50μM bicuculline (Tocris Israel,) and tetrodotoxin (TTX) 1 μM (Tocris) were added to the external ACSF solution. To confirm excitatory currents, cyanquixaline (CNQX) 20 μM (Sigma-Aldrich), D-AP-5 50μM (Tocris) were added to the ASCF.

To record mIPSCs, the internal solution contained (in mM): 140 cesium chloride, 1 EGTA, 6 KCl, 4 NaCl, 2 MgCl2, and 10 HEPES, pH 7.25 and 280 mOsm. In these conditions, the reversal potential of chloride is ∼0 mV. CNQX 20 μM (Sigma-Aldrich), D-AP-550μM (Tocris), and TTX 1 μM (Tocris) were also added to the external ACSF solution. mIPSCs were recorded in voltage clamp mode at a holding potential of −60 mV. To confirm inhibitory currents, 50-μM bicuculline (Tocris) was added to the bath to block GABA_A_ receptors. Data were acquired by Double IPA® Integrated Patch Clamp Amplifiers with Data Acquisition System (Sutter Instruments®). Data was sampled at 20 kHz and filtered at 2 kHz. Series resistance, Rin, and membrane capacitance were monitored throughout, and experiments where resistance changed, was >20% were discarded. Data quantification was done with Sutter Patch (Version 2.2, Sutter Instruments, Novato, CA) and Clampfit (Molecular Devices, Sunnyvale, CA), and subsequently analyzed using GraphPad Prism®.

### Verification of chemogenetic pyramidal neuron activity inhibition

Electrophysiological validation of the activity manipulation of PrL neurons was conducted as previously described (Kayyal et al., 2019). Briefly, PrL slices were prepared as described above. Pyramidal neurons expressing DREADDs or control viruses were identified by mCherry fluorescence. Whole-cell current clamp recordings were made from the soma of PrL LV pyramidal neurons. After 3–5min stable baseline recording, CNO (10μm) was added to the ASCF solution containing CNQX (20μm), APV (50μm), bicuculline methiodide (20μm), and CAS40709–69-1 (Tocris Bioscience); and applied through the bath to the slice to isolate postsynaptic effects. The changes in resting membrane potential were measured 1-2 min after CNO application. Data quantification was done with Sutter Patch (Version 2.2, Sutter Instruments, Novato, CA) and subsequently analyzed using GraphPad Prism®.

### Behavioral tests

Behavioral tests were conducted as previously described (Bauminger et al., 2022). All rats were tested in all behavioral procedures in the same order. Behavioral tests were conducted under dim lighting (15–20 lx). The battery of tests included the OF test, the NOR task and the SIT. In this study we were interested in the effects of chemogenetic attenuation of pyramidal cells on cognitive function and social behavior, therefore CNO was administered before the NOR task and all rats were consecutively tested in the SIT (i.e., while still under the influence of CNO).

Briefly, the OF test was used to examine general locomotor function (total distance, cm, divided into 5 min bins) and novelty-induced anxiogenic behavior (time in arena center, first 5 min and all 30 min of the test). The NOR task, with an inter-trial interval (ITI) of 5 min, was used to test novelty recognition and working memory performance. We assessed total exploration time (s) and the mean discrimination index (DI), calculated as 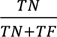 (TN=time of novel object exploration, TF=time of familiar object exploration) during the test phase. The SIT was used to examine social behaviors (e.g., sniffing, physical touch, climbing) and non-social behaviors (e.g., self-grooming, remaining alone), and a sociability index was calculated as the time spent engaging in social behaviors divided by the total test time (5 min).

### Stereotactic surgery for chemogenetic viral injections

Stereotactic surgeries were performed on male rats aged 60-68 PND. Rats were anesthetized with Domitor (2%, 10 mg/kg, subcutaneous) and Ketamine (10%, 100 mg/kg, subcutaneous) (Vetmarket, Modiin, Israel) and treated with Norocarp (1%, 50mg/ml). A total of 1µL of the pAAV-CaMKIIa-hM4D(Gi)-mCherry viral vector (Addgene #50477) or the control vector (pAAV-CaMKIIa-mCherry; Addgene #114469) were bilaterally infused using a Hamilton micro-syringe (Stoelting, Wood Dale, IL, USA) into PrL subregion of mPFC (coordinates: A/P +3.24; M/L ±0.7; V −3.4) at a rate of 0.1µL/min. The needle was held in place for 5 additional minutes before being slowly withdrawn. Animals were allowed 4 weeks of recovery and viral expression time before initiation of behavioral tests.

### Perfusion and histology

A week after the behavioral tests, rats were anesthetized and sacrificed for histological validation of viral injection and expression. Rats were transcardially perfused with PBS, followed by a 4% formaldehyde solution (PFA, Sigma-Aldrich). Brains were fixated overnight and were then transferred to 30% sucrose solution for 48h and next kept at −80 before being sectioned in the cryostat. 45µm sections were collected, washed in PBS, and mounted on glass slides using PBS as a mounting solution, and left to dry for 24h. Slides were visualized using a confocal microscope at 5× 10× and 40× zoom (ZEISS, Jena, Germany).

### Statistical analysis

Results are expressed as means±SEM. For behavioral analysis, two-way ANOVAs and t-tests were used as indicated. Post hoc comparisons were made using unpaired t-tests. Data were analyzed using SPSS27 (IBM, Chicago, IL, USA) and GraphPad Prism®. Normality assumption was examined using the Kolmogorov–Smirnov and Shapiro–Wilk tests, D’Agostino & Pearson tests. For electrophysiological studies, the intrinsic properties were determined with two-tailed unpaired, paired t-tests, and two-way ANOVA followed by Šídák’s multiple comparisons test. Mann Whitney test and Wilcoxon matched pairs signed rank test were used for non-normal distributed data in electrophysiology experiment with DREADD’s+CNO. For all tests significance was determined as ∗p < 0.05.

## Results

### Experiment 1: long-term electrophysiological effects of early-adolescence MK-801

Synaptic properties: Adolescent MK-801 treatment reduced the frequency of mIPSCs onto PrL LV pyramidal neurons.

To test whether adolescent NMDA receptor blockade would impact PrL pyramidal cells activity in adulthood, we conducted a whole-cell voltage-clamp experiment, recording from LV pyramidal neurons in PrL slices. First, we investigated the effects of early-adolescence MK-801 administration on excitatory postsynaptic currents (see representative traces of MK-801 (middle)- and saline (top)-treated rats in Figure 2A; the glutamate receptor blockers CNQX and APV suppressed both the frequency and amplitude of these events (2A, bottom). No group differences were observed in the frequencies of mEPSCs (2B; Saline: 3.610±0.6534 Hz, n=14 cells from 4 Rats; MK-801: 3.679±0.6267 Hz, n=12 cells from 4 Rats; P=0.9704) and amplitude (2C; saline: 14.25±0.9085; MK-801: 13.81±0.8005; p=0.7424.

**Figure 2.**
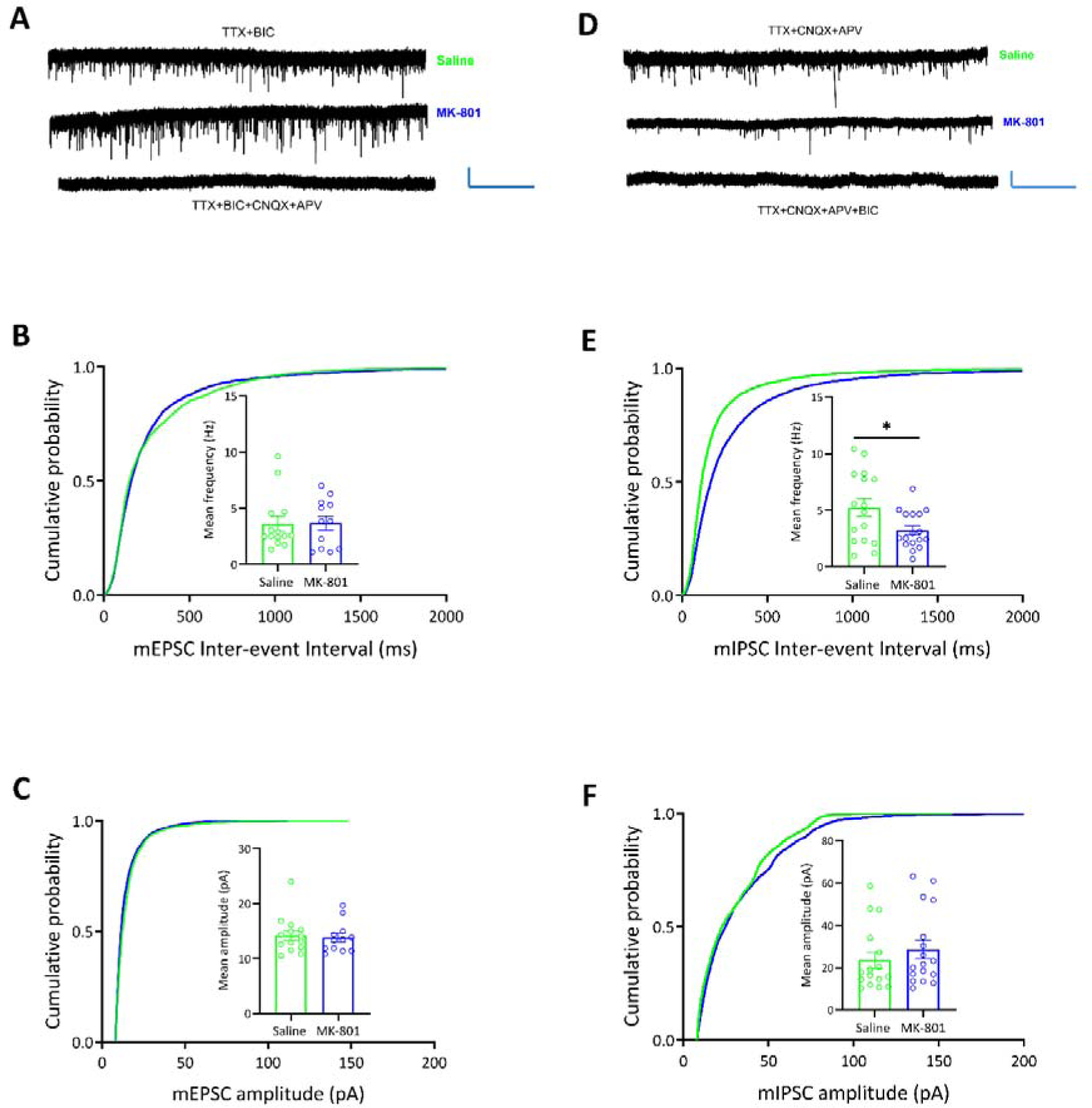
Effects of early-adolescence MK-801 administration on excitatory and inhibitory synaptic transmission in adulthood. (A) Representative traces of mEPSCs for each treatment; Scale bars 20 mV and 10 Seconds. (B) No difference was observed in mEPSCs frequencies between early-adolescence MK-801 treated-rats and saline controls. (C) No difference was observed in mEPSCs amplitudes between early-adolescence MK-801 treated-rats and saline controls. (D) Representative traces of mIPSCs for each treatment; Scale bars 20 mV and 10 Seconds. (E) Early-adolescence chronic MK-801 administration reduced the mIPSCs frequencies compared to saline controls. (F) No difference was observed in mIPSCs amplitudes between early-adolescence MK-801 treated rats and saline controls. \**p*<0.05.

Next, to test the effects of early-adolescence MK-801 administration on inhibitory postsynaptic currents, we pharmacologically inhibited all excitatory synaptic input by isolating the GABAergic currents and recorded mIPSCs from PrL pyramidal neurons, see representative traces of MK-801 (middle)- and saline (top)-treated rats (2D); the GABA_A_ receptor antagonist bicuculline suppressed both the frequency and amplitude of these events (2D, bottom). The frequencies of mIPSCs onto PrL LV pyramidal neurons from brain slices obtained from early-adolescence MK-801-treated rats (n=17 cells from 4 rats) were significantly reduced compared to saline controls (n=16 cells from 5 rats) (2E; *p*=0.0266). There were no group differences in the amplitude of mIPSCs (2F, *p*=0.3823). Consistent with previous reports (Flores-Barrera et al., 2020), these results indicate that NMDA receptor blockade in early adolescence induces lasting impairments in presynaptic inhibitory release in adulthood but does not lead to long-term changes in excitatory presynaptic activity.

Intrinsic properties: Adolescent MK-801 treatment does not change the intrinsic properties of PrL LV pyramidal neurons.

To examine whether early-adolescence MK-801 chronic treatment modifies the intrinsic properties of LV PrL pyramidal neurons, we performed a whole cell current clamp experiment (see extended electrophysiological data in Figure 2-1). No group differences were found in PrL LV pyramidal neurons firing frequencies (F_(8, 208)_=0.2275, p=0.9856; 2-1A, B) and other intrinsic properties like RMP, mAHP, sAHP, Input resistance, sag ratios, time constants, AP amplitude, AP half-width, AP threshold, and rheobase currents (2-1C-L; Table 1). This indicates that early-adolescence MK-801 administration induces hypofunction of GABAergic synaptic release during adulthood, without altering glutamatergic synaptic and intrinsic properties of PrL LV pyramidal neurons.

**Figure 2-1.**
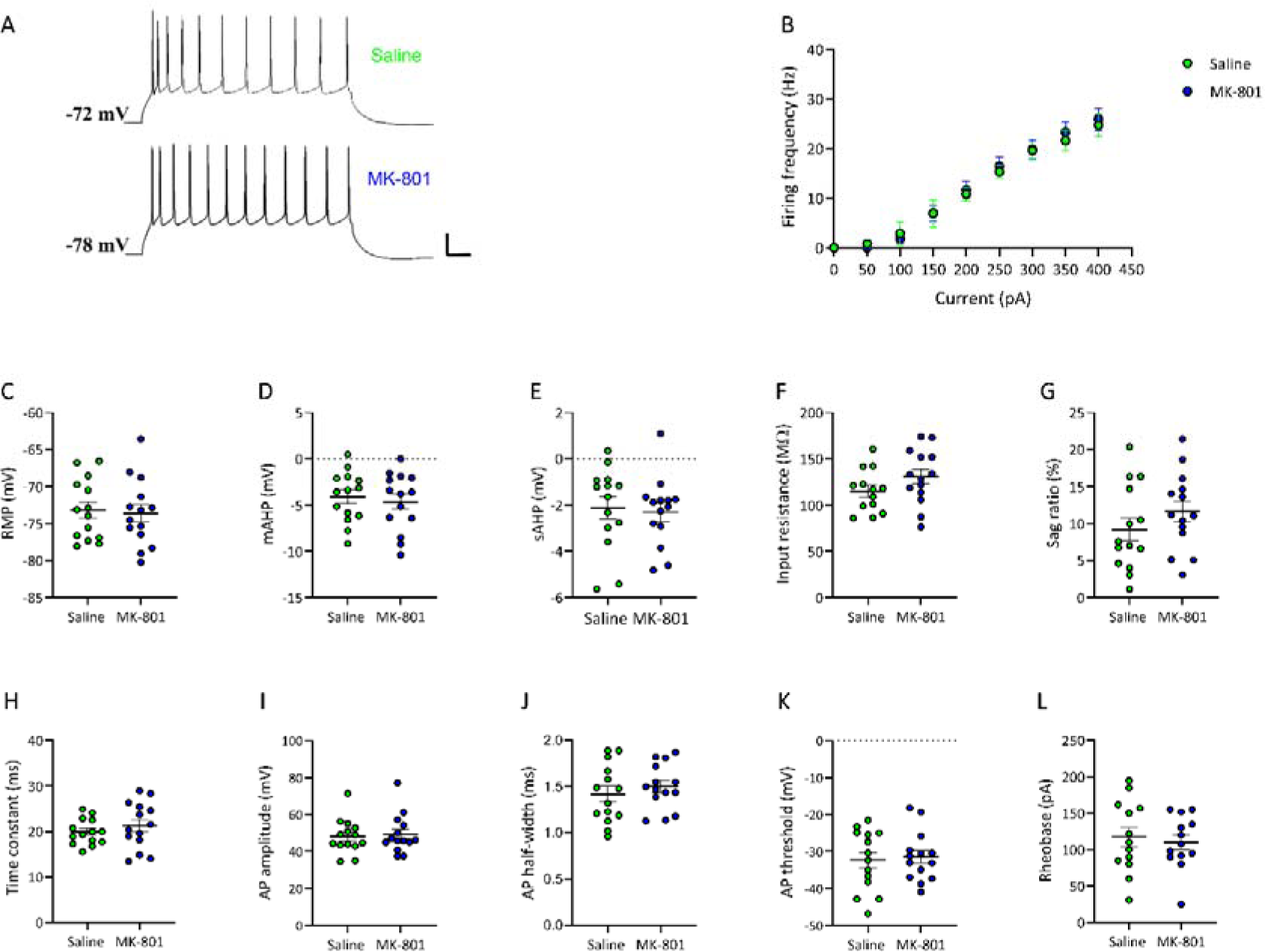
Effects of early-adolescence MK-801 administration on intrinsic properties of PrL LV pyramidal cells in adulthood. (A) Representative traces of firing frequencies following early-adolescence saline (upper) or MK-801 (lower) treatments, scale bar 20 mV and 50 ms from 350 pA step current injections. No differences were found between early-adolescent saline or MK-801 treatments in: (B) firing frequencies, (C) RMP, (D) mAHP, (E) sAHP, (F) Input resistance, (G) sag ratios, (H) time constants, (I) AP amplitude, (J) AP half-width, (K) AP threshold, and (L) rheobase currents.

**Table 1.**
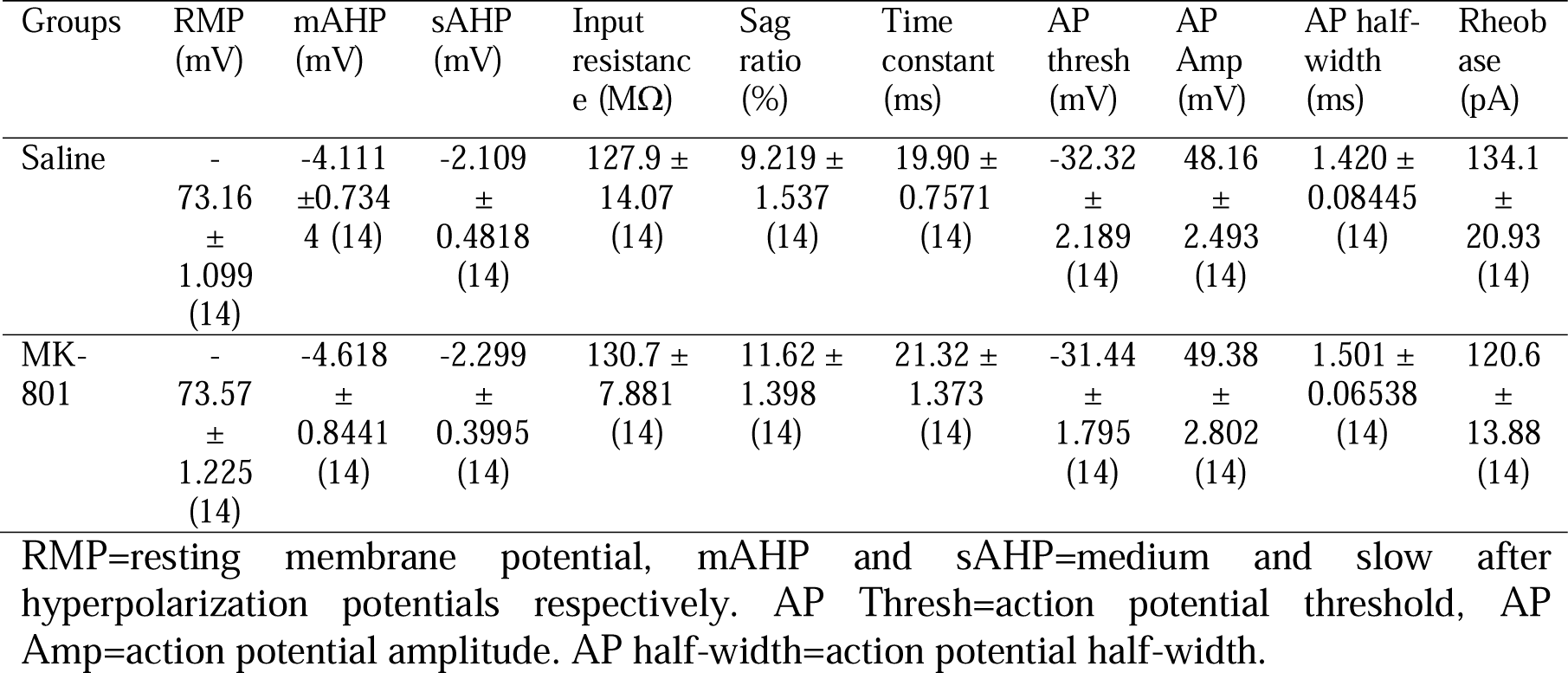
Values are expressed as mean±SEM. The number of cells is in parentheses.

### Experiment 2 – Behavioral effects of chemogenetic attenuation of PrL pyramidal neuron activity in MK-801 or saline-treated rats

Since our electrophysiological findings point to reduced inhibitory drive, i.e. an enhanced E/I balance, we asked whether normalizing this imbalance by reducing pyramidal neuron activity would reverse the behavioral deficits that were previously observed in rats administered with MK-801 in early adolescence (Bauminger et al., 2022).

Viral validation: Rats were injected with viral constructs encoding hM4Di in CaMKII-expressing neurons. As can be seen in Figure 3A, hM4Di-coupled mCherry fluorescence was observed within a well-defined region within the PrL mPFC. Next, using whole cell patch clamp electrophysiology (3B), we validated the inhibitory effect of CNO in PrL brain slices of rats injected with a hM4Di or control virus. Figure 3C shows representative traces of CNO (10uM) effect on cells infected with control or hM4Di viruses. While the bath application of CNO (10um) on PrL slices did not change the RMP in cells infected with the control virus (3D left; p=0.9681; n=12 cells from 3 rats), a rapid hyperpolarization in RMP of hM4Di-expressing neurons was observed following CNO application (3D right; p=0.0010; n=11 cells from 7 rats).

**Figure 3.**
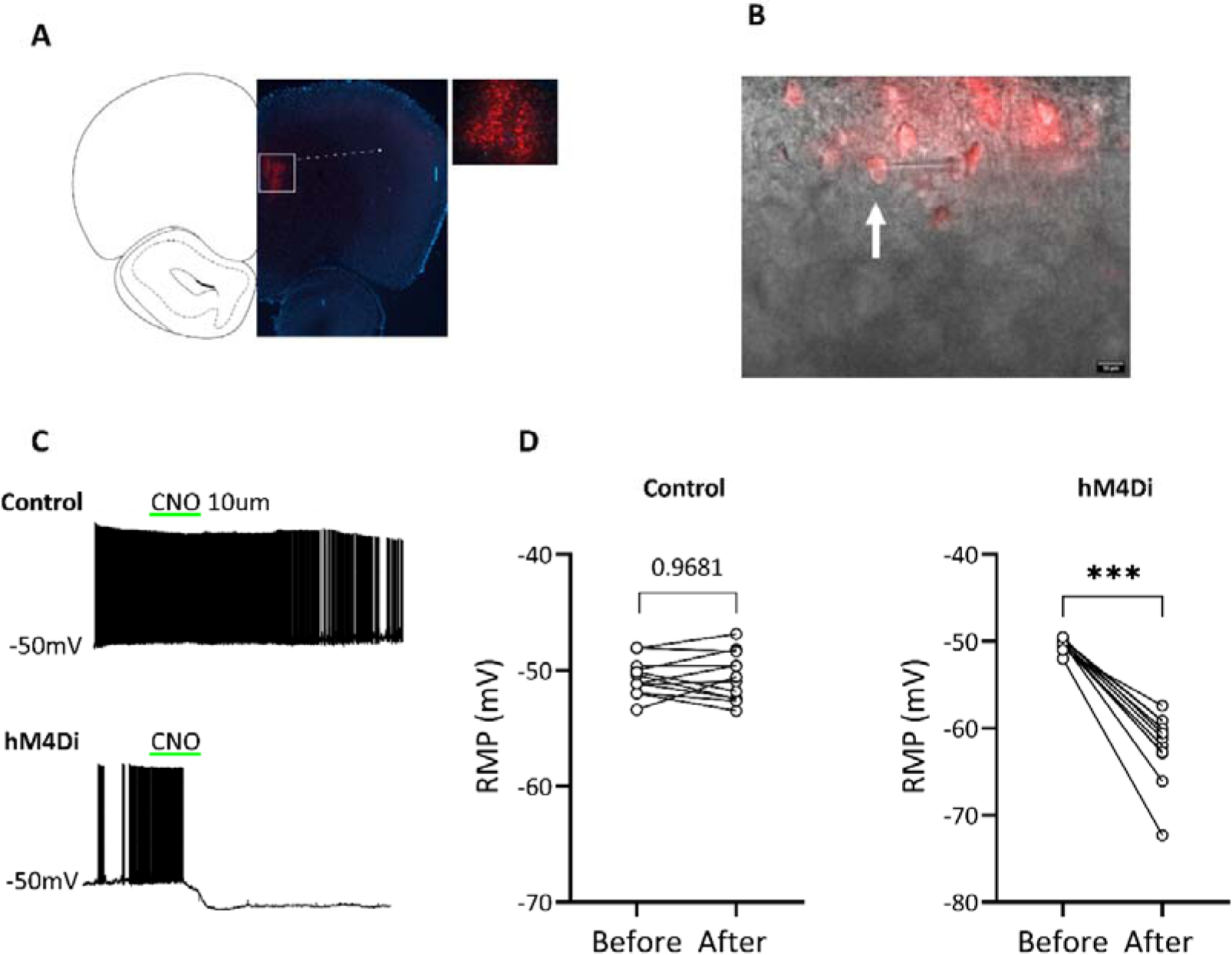
Validation of viral expression and activity. (A) Representative photomicrographs of viral-mediated gene transfer showing hM4Di-coupled mCherry fluorescence in PrL. (B) Representative photomicrographs of electrophysiological recordings from mCherry expressing neuron, the arrow indicates the patch pipette position for recorded neuron. (C) Representative traces showing the effects of CNO on hM4Di (lower) and control virus (upper) expressing cells, scale bar 20mV and 100 seconds. (D) No change in RMP was observed in cells infected with the control virus following CNO application (left); RMP was reduced following CNO application in hM4Di expressing cells (right). \*\*\**p*<0.001.

Behavioral results: First, to examine baseline locomotor activity and anxiety-like behavior in MK-801-and saline-treated rats, the OF test was performed. An unpaired t-test revealed no difference in total distance traveled (cm) between saline (5122±1079, n=19) or MK-801 (5580±963, n=20) treated rats (Figure 4A left; t(37)=1.399, p=0.170), and no difference in time spent in the center of the arena in the first 5min bin (4A right; saline: 68.6±6.2 n=19, MK-801: 66±7.9 n=20; t(37)=-.25, p=0.8) or time spent in the center of the arena in all 30 min of the open field test (data not shown; saline: 309.8±232.7 n=19, MK-801: 321.3±212 n=20; t(37)=.03, p=.87). These findings confirm our previous findings (Bauminger et al., 2022) of no locomotor deficits or anxiety-like behavior in adult rats treated with MK-801 in early adolescence.

**Figure 4.**
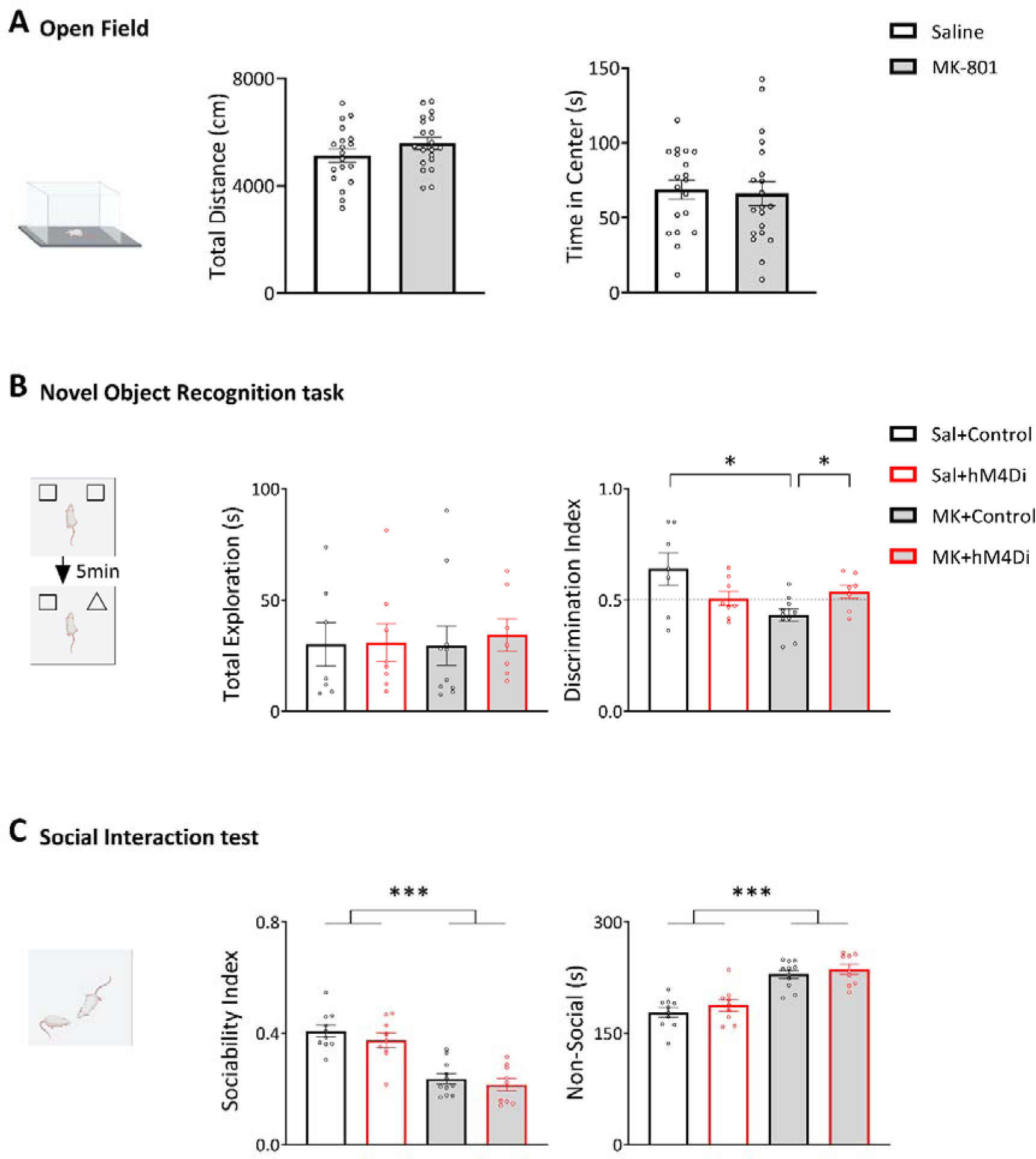
Reversal of early-adolescence MK-801-induced NOR deficits by chemogenetic attenuation of PrL pyramidal neurons firing. (A) In the OF test, early-adolescence MK-801 administration did not impact locomotor behavior (left), or time spent in the center during the first 5min bin (right). (B) In the test phase of the NOR task, no group differences were observed in total exploration time (left) but the MK-801/Control group displayed reduced DI compared to the Saline/Control and MK-801/hM4Di groups (right). (C) In the SIT, MK-801/Control and MK-801/hM4Di showed a reduced Sociability Index (left) and increased non-social behavior (right) compared to the Saline/Control and Saline/hM4Di groups. Bars represent group means and SEM. \**p*<0.05, \*\*\**p*<0.001.

Next, to examine whether chemogenetic attenuation of PrL pyramidal neurons reverses the early-adolescence MK-801-induced recognition memory deficits, the NOR task was performed 30min after CNO administration. CNO was administered to all animals in the experiment. A two-way ANOVA (Drug×Virus, 2×2) on the DI revealed a significant Drug effect (4B right; F(1,28)=4.45, p=0.04) and a significant Drug×Virus interaction (F(1,28)=8, p=.009). Post hoc analysis revealed a reduced DI in the MK-801/Control virus group (.43±.08) compared to the saline/Control virus group (.63±.19; p=0.031) and the MK-801/hM4Di group (.53+.08; p=0.022). No differences were found between the saline/Control virus and MK-801/hM4Di groups (p=0.38) or between the saline/Control virus and saline/hM4Di (p=0.106). There were no group differences in total exploration time (4B left; F(1,28)=0.027, p=0.871). These findings confirm our previous findings of recognition memory deficits in adult rats treated with MK-801 in early adolescence (Bauminger et al., 2022) and show that chemogenetic attenuation of PrL pyramidal neurons reverses this behavioral abnormality without affecting NOR behavior on its own.

In previous studies we found impaired social behavior in early-adolescence MK-801-treated rats (Bauminger et al., 2022). Here, we examined whether chemogenetic attenuation of PrL pyramidal neuron activity can reverse social behavior deficits. Two-way ANOVA’s (Drug×Virus, 2×2) on the sociability index (Figure 4C left) and the time spent engaging in non-social behavior (4C right) revealed significant drug effects in both measures (F(1,35)=56.29, p=0.000; F(1,35)=56.29, p=0.000, respectively), but no Drug×Virus interactions (F(1,35)=0.079, p=.78; F(1,35)=0.079, p=.78). MK-801-treated rats showed a reduced sociability index (0.22±.06) and increased non-social behavior (232±18.92) compared to saline treated rats (sociability index: 0.39±0.07, non-social behavior: 182.31±22.08). These findings replicate our previous findings of social behavior deficits in adult rats treated with MK-801 in early adolescence, and indicate that chemogenetic attenuation of PrL pyramidal neurons does not mitigate this behavioral deficit or produce social behavior abnormalities on its own.

## Discussion

The present findings show that chronic administration of MK-801 during early adolescence leads to long-term deficits in recognition memory and social behavior, and impairs inhibitory activity in the PrL mPFC of adult male rats. Chemogenetic attenuation of pyramidal cells activity in the PrL rescues early-adolescence MK-801-induced deficits in recognition memory, but not social behavior.

Our behavioral findings, i.e., impaired recognition memory and social behavior following early-adolescence NMDAr blockade, replicate our previous findings with this rodent model of SZ-like social and cognitive deficits (Bauminger et al., 2022). The neurodevelopmental approach used in this model highlights the crucial impact of glutamate signaling during adolescence on PFC maturation and, by extension, on PFC-dependent behavior.

Our electrophysiological findings of reduced mIPSCs in adolescent MK-801-treated rats suggest perturbed E/I balance in adulthood following early-adolescence NMDAr hypofunction. Previous studies indicate that acute NMDAr hypofunction leads to excess glutamate release (Moghaddam et al., 1997). In the present study, chronic NMDAr hypofunction during early adolescence did not alter mEPSPs in adulthood. Presumably, acute NMDAr blockade results in excess glutamate release, which is normalized over time, but long-term impairments in GABA interneuron function remain.

Among the discrete GABAergic sub-types of interneurons, parvalbumin (PV)-expressing interneurons are important regulators of pyramidal cell activity. Inhibitory inputs from PV interneurons to pyramidal cells are required for synchronized cortical gamma oscillations (Lewis, Curley, Glausier, & Volk, 2012), a neural mechanism that underlies intact cognitive function (Gonzalez-Burgos & Lewis, 2008). We have shown previously that early-adolescence MK-801 administration reduces the mRNA expression levels of PV in the PrL (Bauminger et al., 2022), and reduced PV protein levels were also found following chronic NMDAr blockade in adolescence (Li et al., 2016) and adulthood (McKibben, Jenkins, Adams, Harte, & Reynolds, 2010). Furthermore, reduced inhibitory activity and impairments in the activity and expression of PV interneuron are hallmark findings in SZ (Dienel & Lewis, 2019; Kaar, Angelescu, Marques, & Howes, 2019). NMDAr hypofunction may induce impairments in this subset of GABAergic interneurons, weakening their inhibitory control over pyramidal cells activity, creating a dysregulated E/I balance and consequently leading to the emergence of cognitive deficits.

The reversal of early-adolescence MK-801-induced recognition memory deficits by chemogenetic attenuation of pyramidal cells in the PrL mPFC suggests that restored E/I balance has therapeutic effects on cognitive abnormalities. In line with this, Levetiracetam is a promising SZ drug candidate which restores E/I balance abnormalities via its actions on KV3.1 channels on fast-spiking PV interneurons (Huang, Tsai, Huang, & Wu, 2009). This drug was found in preclinical studies to reverse NOR deficits emerging following chronic adolescent stress (Cavichioli, Santos-Silva, Grace, Guimarães, & Gomes, 2022), and ketamine-induced memory impairments (Koh, Shao, Rosenzweig-Lipson, & Gallagher, 2018). Furthermore, in a randomized controlled clinical trial, levetiracetam was found efficient in improving cognitive function among patients with SZ (Behdani et al., 2022). It is plausible that attenuated excitability of pyramidal cells mimicked the control of PV INs on pyramidal cells and thus led to positive outcomes on cognitive behavior. In support of this, chemogenetic activation of PV interneurons was found to reverse cognitive deficits induced by adolescent MK-801 (Chamberlin et al., 2023; Huang et al., 2021). Hence, both reduced excitation and increased inhibition might lead to beneficial outcomes on cognitive impairments. Future drug development efforts should focus on reducing the excitatory drive, rather than or in addition to increasing the inhibitory drive, as a therapeutic outcome.

Chemogenetic attenuation of PrL pyramidal cells did not reverse the early-adolescence MK-801-induced deficits in social behavior. Previously we have shown that endocannabinoid enhancement reversed the early-adolescence MK-801-induced cognitive deficits via cannabinoid receptor 2 activation, and the social deficits via cannabinoid receptor 1 activation (Bauminger et al., 2022), highlighting that social and cognitive behaviors rely on different neural mechanisms. Social behavior abnormalities in the current model may be related to other circuits, such as the ventral tegmental area-nucleus accumbens pathway (Gunaydin et al., 2014) or the basolateral amygdala-anterior cingulate cortex pathway (Huang et al., 2022). Future studies should examine the ability to reverse the early-adolescence MK-801-induced social behavior abnormalities using chemogenetic tools in other cell types or neural circuits.

To the best of our knowledge, this study is the first to examine the ability of attenuated PrL pyramidal cell activity to reverse the early-adolescence MK-801-induced SZ-like behavioral abnormalities. Our findings have important implications on the understanding of the long-term effects of NMDAr blockade during adolescence. Furthermore, these findings highlight the importance of focusing on restoring the E/I balance in future drug development to address cognitive dysfunction in SZ and other psychiatric and neurological illnesses.

## Acknowledgments

This study was funded by Israel Ministry of Health (MOH) grant 3-15065 to IGS and IA, and by Israel Science Foundation (ISF) grant 1481/20 to IGS. We thank Professor Kobi Rosenblum for providing the electrophysiological tools and Dr. Tomer Mizrachi Zer-Aviv for his help with perfusion.

